# CoPhosK: A Method for Comprehensive Kinase Substrate Annotation Using Co-phosphorylation Analysis

**DOI:** 10.1101/251009

**Authors:** Marzieh Ayati, Danica Wiredja, Daniela Schlatzer, Sean Maxwell, Ming Li, Mehmet Koyutürk, Mark R. Chance

## Abstract

We present CoPhosK to predict kinase-substrate associations for phosphopeptide substrates detected by mass spectrometry (MS). The tool utilizes a Naïve Bayes framework with priors of known kinase-substrate associations (KSAs) to generate its predictions. Through the mining of MS data for the collective dynamic signatures of the kinases’ substrates revealed by correlation analysis of phosphopeptide intensity data, the tool infers KSAs in the data for the considerable body of substrates lacking such annotations. We benchmarked the tool against existing approaches for predicting KSAs that rely on static information (e.g. sequences, structures and interactions) using publically available MS data, including breast, colon, and ovarian cancer models. The benchmarking reveals that co-phosphorylation analysis can significantly improve prediction performance when static information is available (about 35% of sites) while providing reliable predictions for the remainder, thus tripling the KSAs available from the experimental MS data providing a to comprehensive and reliable characterization of the landscape of kinase-substrate interactions well beyond current limitations.

**Author Summary:** Kinases play an important role in cellular regulation and have emerged as an important class of drug targets for many diseases, particularly cancers. Comprehensive identification of the links between kinases and their substrates enhances our ability to understand the underlying mechanism of diseases and signalling networks to drive drug discovery. Most of the current computational methods for prediction of kinase-substrate associations use static information such as sequence motifs and physical interactions to generate predictions. However, phosphorylation is a dynamic process and these static predictions may overlook unique features of cellular context, where kinases may be rewired. In this manuscript, we propose a computational method, CoPhosK, which uses the mass spectrometry based phosphoproteomics data to predict the kinase for all identified phosphosites in the experiment. We show that our approach complements and extends existing approaches.

## Introduction

Protein phosphorylation (PP) is a post-translational modification that is central to cellular signalling where networks composed of kinases, phosphatases, and their substrates regulate the sites and levels of phosphorylation at the molecular level. MS-based approaches based on phosphopeptide enrichment can report the identity and intensity of thousands of protein phosphorylation sites in the context of their phosphopeptides [1-4] and these data have populated several public databases of phosphosites (e.g. phospho.ELM [5] and PhosphoSitePlus [6]) with over 500,000 sites already identified in humans. Despite this success in identifying cellular kinase substrates, the identity of the responsible kinase is typically unknown for 90% of the cases. For this reason, effective bioinformatics tools to predict KSAs have been essential to filling the gap between phosphosite identification and kinase annotation.

Prior KSA prediction methods have focused mainly on the consensus sequence motifs recognized by the active sites of kinases [7-11]. The modest specificity of these methods led researchers to integrate other contextual information such as protein structure and physical interactions between proteins [12,13]. Addition of these cues to the analysis has enhanced the accuracy of prediction methods [14]. The latest improvements have reduced hub bias in the predictions [15] and are promulgated in the software KinomeXplorer (Figure 1). Weaknesses of these approaches include: 1) the network predictions are static and 2) incomplete information on protein structure and protein interactions permit predictions of KSAs for only 30-40% of the sites observed in MS experiments. As cellular signalling is a highly dynamic and transient process, the incorporation of specific experimental information on the dynamics of phosphorylation (e.g. the changes in phosphosite occupancy and intensity across selected biological states) into KSA predictions could provide a key to permitting “global” scale predictions.

**Figure 1.**
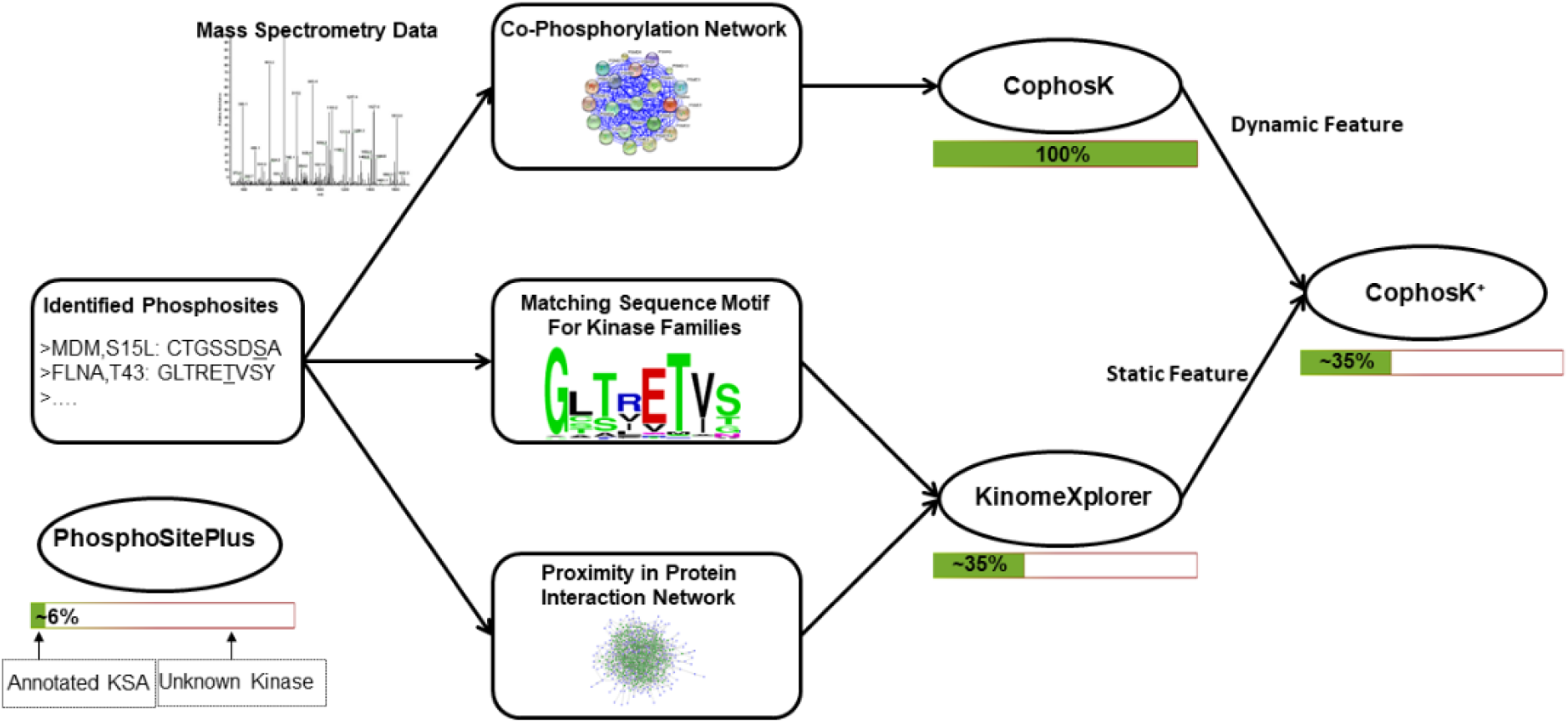
The overview of CophosK and CophosK+. After identification of phosphorylated sites, KinomeXplorer utilizes sequence match scoring and network proximity of kinases and substrates to predict KSAs. CophosK constructs the co-phosphorylation network in order to infer KSAs. CophosK+ combines all the scores to provide more accurate KSA predictions. PhosphoSitePlus annotates ~6% of the identified phosphosites by their associated kinases. KinomeXplorer improves the coverage of annotations to 35% and can improve accuracy of predictions in the context of co-phosphorylation information (CoPhosK+). On the other hand, CophosK alone is able to annotate 100% of the identified phosphosites in the experiment.

The payoff is that identifying the kinases active with respect to specific substrates can provide critical insights into biological processes in both development and disease. In particular, kinase inhibition strategies for combating multiple cancer related phenotypes have leveraged an understanding of the phosphorylation dynamics of cellular systems through elucidation of kinase-substrate networks [16-18]. As of early 2018, over 40 kinase inhibitors have received US FDA approval for the treatment of various diseases, most of them related to cancer [19]. Overall, a detailed understanding of phosphorylation networks is critical for drug discovery and development as well as understanding the basic biology of signalling. To address this challenge, we present CophosK, an algorithmic pipeline that extracts *co-phosphorylation* patterns from MS data to provide global predictions of KSAs (Figure 1). Our results show that pairs of phosphosites that are substrates of the same kinase are significantly more likely to be co-phosphorylated (i.e., exhibit correlated phosphorylation levels across different biological states and/or over time), as compared to other pairs of phosphosites. Motivated by this observation, our algorithm for predicting kinase-substrates associations is based on the following principle: If a pair of phosphosites exhibit similar phosphorylation patterns through different biological states, and if we know that a specific kinase acts on one of these phosphosites, it is likely that the kinase acts on the other phosphosite as well. Based on this principle, we develop a Bayesian framework that utilizes multiple co-phosphorylation relationships of a phosphosite to score and rank kinases for that phosphosite.

## Results

### MS based phosphorylation data to probe biological states for co-phosphorylation detection

To develop and validate our method, we initially analyzed two following datasets:

*(I) Breast Cancer (BC) Patient-Derived Xenografts (PDX) tissues*: Huang *et al.* used the proteomic method called isobaric tags for relative and absolute quantification (iTRAQ) to identify 56874 phosphosites in 24 breast cancer PDX models [20]. In this study, we removed phosphosites with missing intensity values in any sample. This resulted in intensity data for 15780 phosphosites from 4539 proteins, where 13840 serines, 2280 threonines and 67 tyrosines are phosphorylated. As suggested by Huang *et al*, the reference sample used to compute fold change is comprised of a standard sample from 16 out of 24 tumours with equal amounts from each tumour.

*(II) Ovarian Cancer (OC) tumors*: The Clinical Proteomic Tumor Analysis Consortium conducted an extensive MS based phosphoproteomic of ovarian HGSC tumors characterized by The Cancer Genome Atlas [21]. They have reported 24429 phosphosites from 6769 phosphoproteins in 69 tumors. We filtered out the phosphosites with missing data and also selected a subset of tumors to cover more phosphosites. This resulted in a total of 5017 phosphosites from 2425 proteins in 12 tumors where 4258 serines, 657 threonines and 102 tyrosines are phosphorylated.

Then we applied the proposed method to predict KSAs and assess the performance of the methods on four other independent datasets:

*(III) BC PDX*: The NCI Clinical Proteomic Tumor Analysis Consortium (CPTAC) conducted an extensive MS based phosphoproteomic of TCGA breast cancer samples [22]. After selecting the subset of samples to have the highest coverage and filtering the phosphosites with missing intensity values in those tumors, the remaining data contained intensity values for 11018 phosphosites mapping to 8304 phosphoproteins in 20 tumors.

*(IV, V) Breast Cancer (BC) PDX and Ovarian Cancer (OC) PDX*: This dataset was generated to analyze the effect of delayed cold Ischemia on the stability of phosphoproteins in tumor samples using quantitative LC-MS/MS. The phosphorylation level of tumor samples have been measured across 3 time points [23]. After phosphosites with missing intensity values were filtered out, the dataset included 4802 phosphosites corresponding to 2230 phosphoproteins in 12 ovarian tumors and 8150 phosphosites mapping to 3025 phosphosproteins in 18 breast cancer xenograft tissues.

*(VI) Colorectal cancer (CC)*: Abe *et al*. performed immobilized metal-ion affinity chromatography-based phosphoproteomic and highly sensitive pY proteomic analyses to obtain deep phosphoproteomic data in 4 different colorectal cancer cell lines [24]. The dataset included 5382 phosphosites with intensity values cross 12 different conditions. These phosphosites mapped to 2230 phosphoproteins.

### Evidence for Statistically Significant Co-phosphorylation Distributions

We used the data above to address two initial questions related to the appropriate development of our phosphorylation analysis method. First, we considered the correlation analysis approach appropriate for these data and second; we considered the effect of experimental design (e.g. number of independent samples). For the latter, we intuited that a larger number of samples should provide greater opportunity to sample a wider range of phosphorylation states based on sampling a wider range of biological variations. For the analysis, we define the vector containing the phosphorylation levels of a phosphosite across a number of biological states as the *phosphorylation profile* of a phosphosite for which co-phosphorylation will be evaluated using fold-change values in this vector. For assessing the correlations of these vectors across the data matrix, we compared the suitability of various mathematical methods from the literature. For example, for gene co-expression, Pearson’s correlation is the most popular correlation measure used to capture linear relations between the expression profiles of genes [25], while mutual information is commonly utilized to describe non-linear relationships [26]. Song *et al.* proposed biweight-midcorrelation as an alternative, and showed that it outperforms mutual information in terms elucidating pairwise relationships between genes; it is also more robust than Pearson correlation with respect to outliers [27]. Therefore, we chose biweight-midcorrelation to assess the correlation between the phosphorylation profiles of phosphosites.

As a second consideration, we reasoned that the number of biological states (i.e., the number of dimensions in the phosphorylation profiles of phosphosites, which is equal to the number of independent samples/tumors in our datasets) may have a significant effect on the interpretation and reliability of our co-phosphorylation analysis, thus we investigated the effect of number of states on the distribution of co-phosphorylation across all pairs of phosphosites. Using the 24 breast cancer PDX samples for testing, we randomly selected 3, 6 and 12 samples and analyzed the co-phosphorylation distribution for phosphorylation profiles composed of these randomly selected samples. Then, we randomized the phosphoproteomics data (see below) and compared the distribution of co-phosphorylation among phosphosites in the original data against the randomized data (S1 Fig). We observed that for both the original data and randomized data, co-phosphorylation is normally distributed with a mean around zero when three or more independent biological states are analyzed, and the standard deviation of this distribution goes down with increasing number of samples (i.e., the correlation between phosphorylation profiles tends to be higher when the number of dimensions is lower). However, as the number of biological states in the study increases, the co-phosphorylation distribution among pairs of phosphosites becomes statistically significant and more easily distinguishable from random data (solid vs. dashed lines in S1 Fig). As seen in S1 Fig(b), the curve is steeply sloping below 5-dimensions and flattens out considerably above 10 dimensions. To provide further guidance for benchmarking the method, we carried out simulations using synthetic data. For this purpose, we generated thousands of random vectors (using normal distributions) by varying number of dimensions and plotted the distribution of correlation among pairs of vectors (S2 Fig). Based on this analysis, at least five biological states (or dimensions) are recommended for capturing statistically significant associations. As our datasets had 12-and 24 dimensions, this analysis indicates they were suitable for further analysis. We also show that selection of different samples with fixed number of dimensions do not have a significant effect on co-phosphorylation distribution (S3 Fig).

We compared the distribution of co-phosphorylation for both ovarian (12-dimensions) and breast cancer (24-dimensions) datasets against three different null models to assess the statistical significance of co-phosphorylation for pairs of phosphosites that were detected in these experiments. These null models are constructed via (i) random permutation of the phosphorylation intensities of phosphosites within each sample (biological state), (ii) random permutation of phosphorylation intensities representing the phosphorylation profiles across biological states for each phosphosite, and (iii) random permutation of all intensity values in the phosphosite-biological state matrix. The results of this analysis for the breast cancer cell line dataset are presented in Figure 2. The distribution of co-phosphorylation among pairs of phosphosites in the original dataset is significantly different as compared to the distribution of co-phosphorylation among pairs of phosphosites in all permuted datasets (Kolmogorov-Smirnov (KS) test p-value << 0.01). Namely, the distribution of co-phosphorylation is wider in the original data as compared to any of the three null models; e.g. there are more phosphosite pairs with high positive or negative correlation.

**Figure 2.**
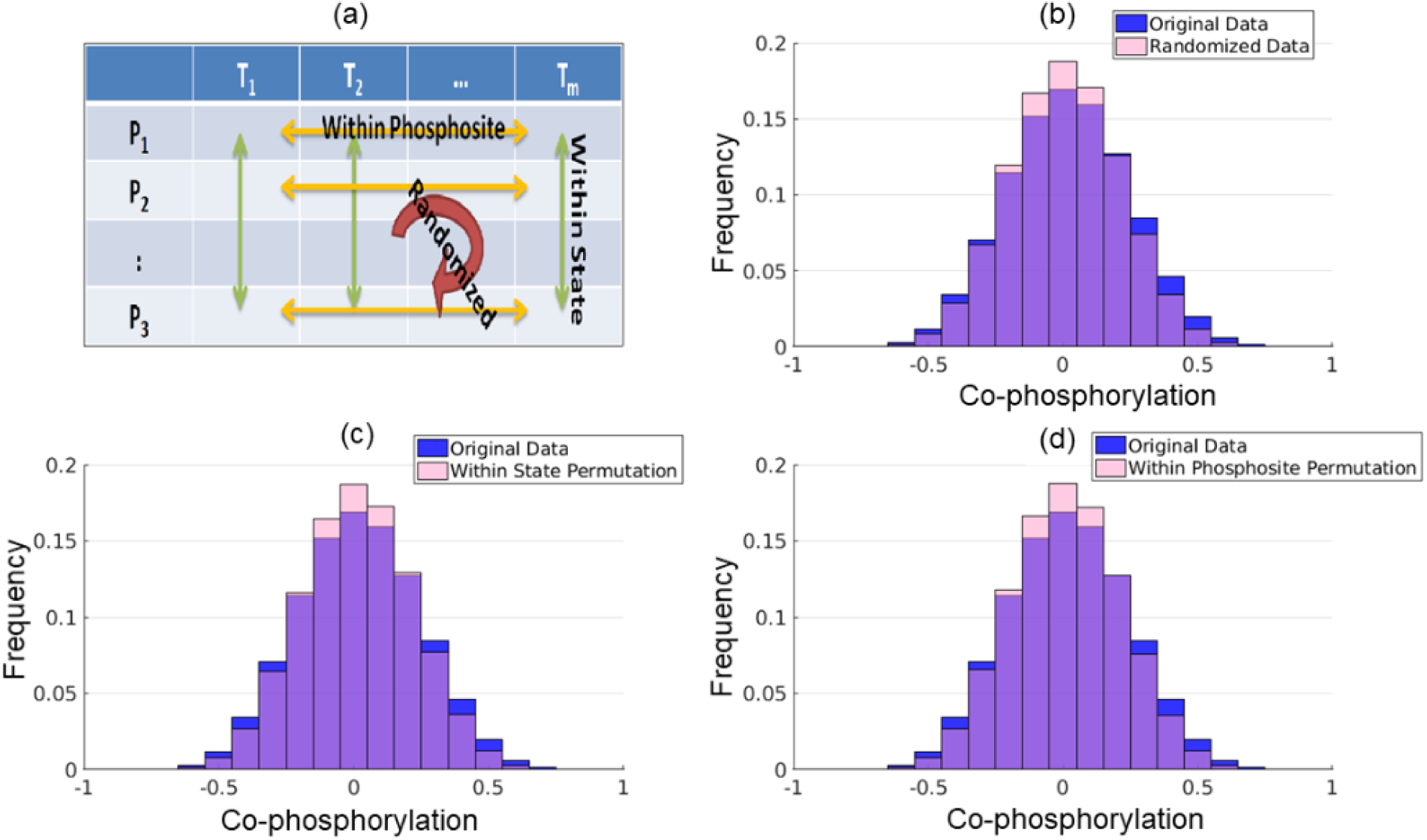
Distribution of co-phosphorylation among pairs of phosphosites on breast cancer PDX. The blue histogram shows the distribution of co-phosphorylation (the correlation between the phosphorylation levels) of all pairs of phosphosites in breast cancer PDX (μ=0.01,σ=0.22). (a) Illustration of the three different permutations tests that were used to assess the significance of this distribution. The pink histogram in each panel shows the distribution of co-phosphorylation of all pairs of phosphosites in 100 permutation representing (b) randomization of all entries in the phosphorylation matrix (μ=0.008,σ=0.20), (c) permutation of all entries across phosphosites for each state (μ=0.01,σ=0.20), and (d) permutation of all entries across states within each phosphosite (μ=0.01,σ=0.20). The distribution of co-phosphorylation in the original dataset is significantly broader as compared to the distribution of co-phosphorylation in all permutations (Kolmogorov-Smirnov (KS) test p-value << 1E-9).

The distribution of co-phosphorylation for ovarian cancer samples is shown in Supplementary S4 Fig; a similar pattern is observed for these data. The statistically significant co-phosphorylation suggests that the correlations may have underlying biological drivers. The same result using Pearson correlation also reported in S5-S7 Fig.

As Li *et al* showed that phosphorylated sites that are modified together tend to participate in similar biological process [28], we hypothesized that phosphosite pairs exhibiting positive correlation of co-phosphorylation may be substrates of the same kinase (e.g. *shared-kinase pairs).* We thus compared the co-phosphorylation distributions (actually sub-distributions) for substrates from the same kinase, as annotated by gold standards, to the original distribution. Using KSAs from PhosphoSitePlus (PSP) for each of the 347 reported kinases, we quantified the co-phosphorylation of all pairs of phosphosites that are listed as that kinase’s substrates (37234 and 8235 shared-kinase pairs in breast cancer and ovarian cancer data, respectively). The distribution of co-phosphorylation of shared-kinase pairs as compared to all other phosphosite pairs is shown in Figure 3. As seen in the figure, these two distributions are significantly different and the co-phosphorylation distribution of shared-kinase pairs is shifted to the right (Kolmogorov-Smirnov test p-value respective < 7.36E-103, 1.1E-6 for breast cancer PDX and ovarian cancer, respectively). In other words, substrates that share a kinase are more likely to be positively co-phosphorylated compared to all pairs of phosphosites.

**Figure 3.**
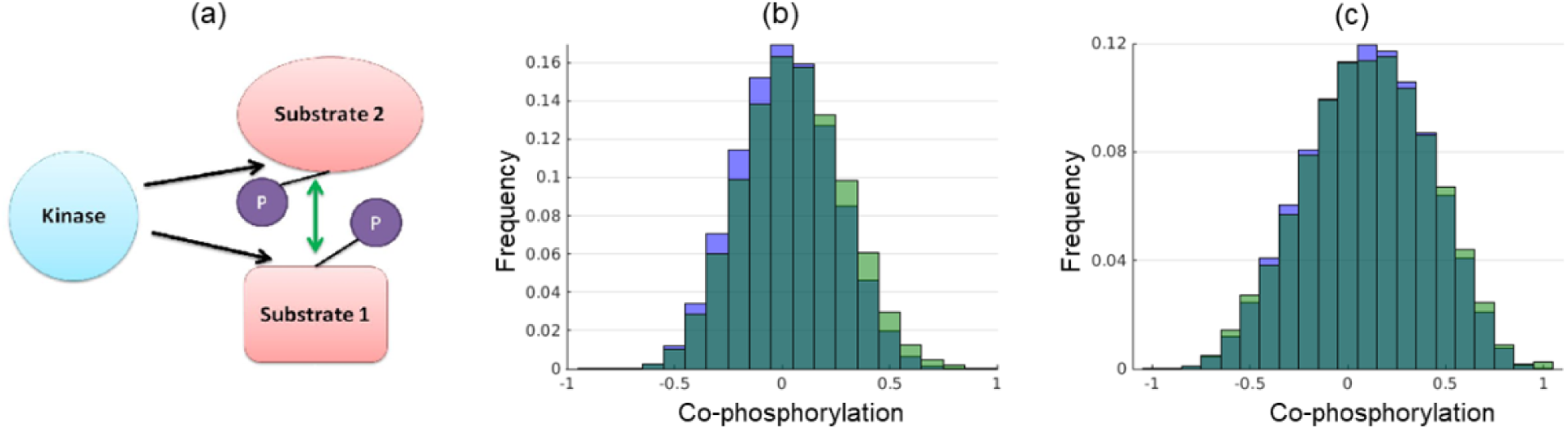
Distribution of co-phosphorylation between phosphosites that are substrates of the same kinase. (a) For each of 347 reported kinases, we compute the co-phosphorylation of all pairs of phosphosites that are reported to be common substrates of that kinase in PhosphositePLUS (*shared-kinase pairs*). In panels (b), (c), the green histogram shows the distribution of co-phosporylation for all *shared-kinase pairs* and the blue histogram shows the distribution of co-phosphorylation for all pairs of phosphosites in the dataset. (b) Breast Cancer PDX dataset (37234 shared kinase pairs; μ=0.05,σ=0.23,kutosis=2.85, skewness=0.10), (c) Ovarian Cancer tumors (8235 shared kinase pairs; μ=0.11,σ=0.32,kutosis=2.54, skewness= −0.10). For both datasets, the distribution for shared-kinase pairs is significantly wider and shifted to the right as compared to the distribution for all phosphosite pairs (KS-test p-value << 1E-9).

Although this positive correlation of substrate pairs can be potentially explained by shared kinase annotation, the data also contains negative correlations of phosphosite pairs (Figure 3) that point to more complex regulatory relationships. One reason for this observation could be that for some phosphosites, multiple kinases have been reported in PhosphoSitePlus, but the annotation does not provide context-specific information, e.g., the kinases might not be active at the same time. The negative co-phosphorylation also might reflect the relationship between substrates and their associated phosphatases since they are expected to follow the opposite pattern in phosphorylation.

### Using Co-Phosphorylation for Kinase-Substrate Association Prediction

Motivated by our observation that substrates of the same kinase are more likely to be positively co-phosphorylated (Figure 3), we developed a co-phosphorylation based prediction method, CophosK, that constructs a co-phosphorylation network (Figure 4) and uses a Naïve Bayes framework on this network to predict KSAs. The idea behind the approach is that the likelihood of the association of a phosphosite with a given kinase is proportional to the fraction of its neighbours in the co-phosphorylation network that are associated with the kinase. This method directly incorporates the dynamics of phosphorylation into KSA prediction methods. Furthermore, since the co-phosphorylation network is context-specific, this method can potentially point to context specific KSAs. Note that this can provide a prediction for all the substrates in the data. Here we are able to score nearly 15,000 KSAs for 101 kinases in breast cancer data and nearly 5,000 KSAs for 75 kinases in the ovarian cancer data. As static information provided by tools like NetworKIN and KinomeXplorer and dynamic information provided by CophosK may be complementary, we also developed CophosK^+^, which is designed to take three main elements into consideration to predict KSAs: Sequence motifs associated with targets of kinases, the network proximity of kinases and substrates in the protein association network, and co-phosphorylation of substrates of kinases (Figure 1). In this case the number of KSA predictions is limited to those that can be predicted by KinomeXplorer.

**Figure 4.**
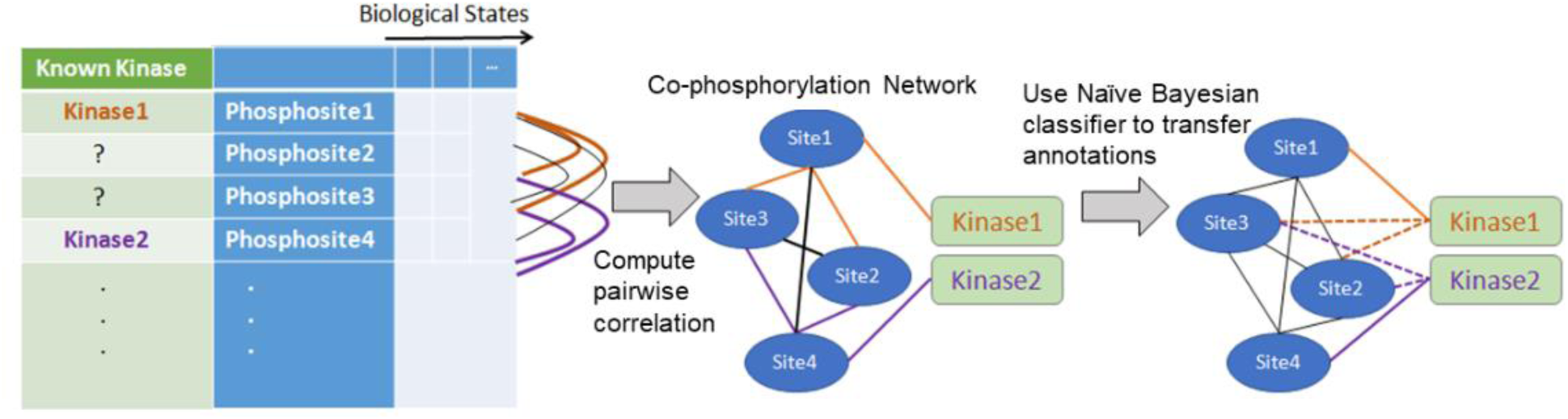
Workflow of CophosK for using co-phosphorylation to predict kinase-substrate associations (KSAs). The method takes as input available information on KSAs and phosphorylation data representing the phosphorylation levels of thousands of phosphosites across multiple biological states. The co-phosphorylation of all pairs of phosphosites is assessed and a co-phosphorylation network is constructed in which phosphosites represent nodes and the weight of the edges is the correlation between phosphosites. CophosK then uses a Naïve Bayes classifier that integrates the interactions in this network with partial information on kinase-substrate interactions to predict new kinase substrate interaction

### Static and dynamic data provide orthogonal KSA predictions

To test the effectiveness of CoPhosK and CophosK^+^, we use leave-one-out cross validation on the list of KSAs reported in PhosphoSitePlus. Namely, for each phosphosite, we hide the association between the phosphosite and its known kinase (called the *target kinase*) and we use other reported KSAs to rank the likely kinases for that phosphosite. To enhance the reliability of the predictions, we only consider kinases that have at least two reported substrates in the database. For each phosphosite in the dataset, we rank all kinases based on the scores computed by CophosK, CoPhosK^+^ and KinomeXplorer and determine the rank of the target kinase.

Figure 5(a) shows the rankings provided by KinomeXplorer (on the y-axis) and the rankings provided by CophosK (on the x-axis) for 313 kinase-substrate association predictions for the ovarian cancer data and 740 kinase-substrate association predictions for the breast cancer PDX data. In the figure, a point that is closer to the origin indicates higher ranking. Figure 5b shows the data in box plot format, this visualization indicates that the distribution of results is overall similar. If the two methods were consistently correct in their predictions, we would see a cluster of points only around the origin. In fact aside the dense cluster around the origin, many predictions are “close to the axes”, indicating a high rank from one approach and a low rank from the competing method. This suggests that these two methods contribute different information, therefore integrating the prediction of these two methods might improve the predictions. Moreover, Figure 5(b) shows the box plot distributions of target kinase rankings for the breast cancer PDX and ovarian cancer data.

**Figure 5.**
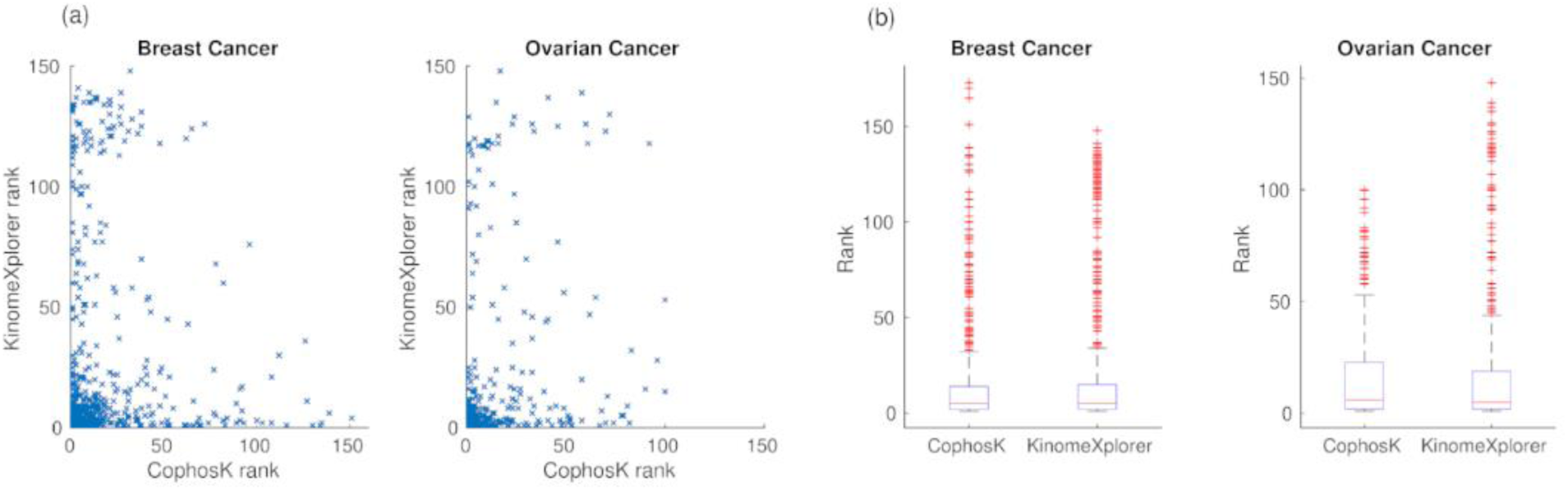
The correspondence between the predictions of CophosK vs. KinomeXplorer in ranking kinases for each phosphosite. For all kinase-substrate associations reported in PhosphoSitePlus for which we can detect a substrate in the LC/MS data, we perform leave-one-out cross validation by hiding the association between the phosphosite and using CophosK to utilize other kinase-substrate associations and co-phosphorylation to rank the likely kinases for the phosphosite. (a) shows the comparison of the rankings provided by CoPhosK (x-axis) against the rankings provided by KinomeXplorer (y-axis) for breast cancer PDX data (740 predictions) and ovarian cancer (313 predictions). In (b), the box plot distribution of the rank of the target kinase according to the prediction of two methods are presented.

To investigate whether CophosK^+^ can exploit any potential synergy between dynamic co-phosphorylation and static predictions, we investigated the performance of CophosK^+^ using the leave-one-out cross validation method described above. Figure 6 shows the overall performance of CophosK, CophosK^+^, KinomeXplorer, and PUEL, an alternative analysis approach [29], when analysed with respect to the six cancer datasets (the residue-specific performance is reported in S8 Fig). In the figure, we report the fraction of phosphosites for which the target kinase is ranked in top 1 and top 5 by each scoring method. As seen in the figure, CophosK and KinomeXplorer deliver similar prediction performance. Since PUEL’s predictions are considerably less accurate on breast cancer (I) and ovarian cancer (II) datasets, we did not run it on other datasets. However, CophosK^+^, our algorithm that comprehensively integrates co-phosphorylation, sequence motifs, and protein interactions, improves the accuracy of KSA predictions over all approaches.

**Figure 6.**
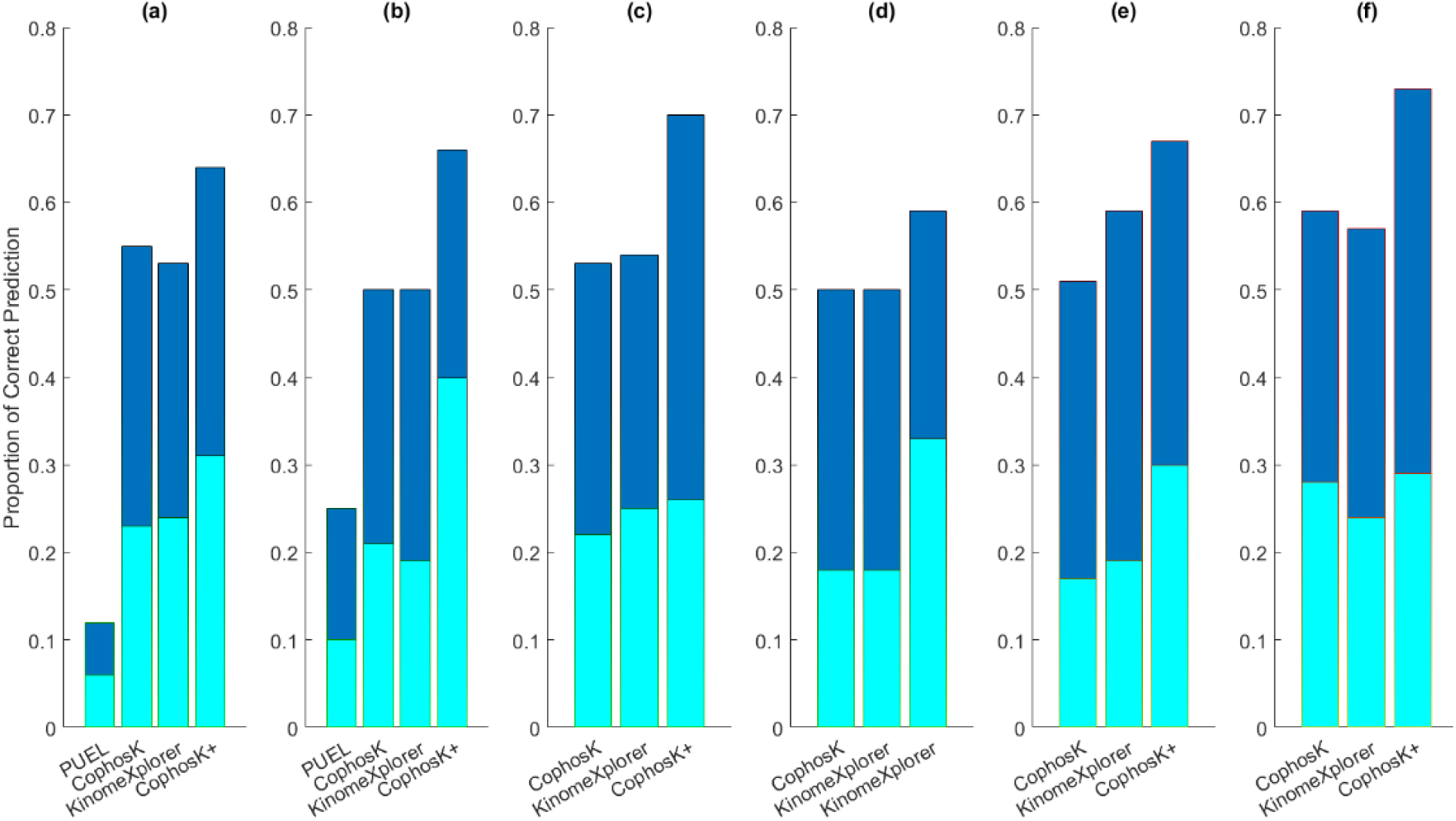
Performance of CophosK, KinomeXplorer and CophosK+ in predicting kinases for phosphosites. For each dataset, we consider all phosphosites that are identified in the dataset and/or reported in PhosphoSitePlus. For each phosphosite, we perform leave-one-out cross validation by hiding the association between the phosphosite and one of its associated kinases (*target kinase*) to rank the likely kinases for the phosphosite using PUEL, CophosK, KinomeXplorer, and CophosK^+^. We report the fraction of phosphosites for which the target kinase is ranked in the top 1 and top 5 predicted kinases by each method (as indicated by different colors in the bar plot). Each panel shows the performance of the methods on (a) breast cancer PDX data(I), (b) ovarian cancer(II), (c) breast cancer(III), (d) breast cancer(IV), (e) ovarian cancer(V), and (f) colorectal cancer(VI) datasets.

### CophosK provides global predictions of KSAs

Next, we investigate the coverage of the phosphoproteome provided by the proposed KSA prediction methods. The number of phosphosites annotated by each method (i.e., the number of phosphosites for which the method was able to make a prediction) is shown in S9 Fig. In the ovarian cancer and breast cancer datasets, respectively 5017 and 15780 phosphosites are identified. Among these, approximately 6% have reported kinase substrate associations in PhosphoSitePlus. If we use KinomeXplorer, we can predict the associated kinase for 47% and 35% of identified phosphosites in ovarian cancer and breast cancer study, respectively.

Since we can compute the co-phosphorylation among all the identified phosphosites in these studies, CoPhosK can predict the associated kinase for all of the phosphosites identified in these studies, providing annotations for 12,000 phosphosites that had no kinase annotation previously available. A downside of these predictions is that their current estimated accuracy is just over 50%, if we consider ranking in the top five a true positive (Figure 6). To this end, CophosK^+^ and CoPhosK together capture the trade-off between adding new annotations and improving existing annotations. Using CophosK^+^ the annotations of phosphosites with existing “static” annotations can be enhanced. Using CoPhosK, on the other hand, new annotations can be also developed for previously uncharacterized phosphoproteins. Thus, the methods create a set of medium confidence predictions for all the phosphosites and high confidence predictions (65%) for the better annotated subset. The kinases predicted for all phosphosites by CophosK and CophosK^+^ are available at compbio.case.edu/cophosk. Any phosphoproteomics data and pre-defined KSAs data can be used for predictions. For example, the results obtained by the application of the proposed methods using phospho.ELM (an alternative phosphosite database) is reported in S10 Fig.

The runtime of CophosK depends on the number of kinases that should be scored for each phosphosite. We assess the runtime of CophosK using a workstation with an AMD Opteron CPU with a 1.9 GHz processor with 64G RAM. For ovarian cancer data, it takes approximately 1 hour and 20 minutes to score 75 kinases for 5017 phosphosites. For breast cancer data, CophosK scores 101 kinases for 15780 phosphosites in approximately 18 hours. Please note that the reporting time is for a single thread process. Since the scoring procedure of each kinase and substrate is independent of each other, this process is highly parallelizable and can be optimized.

### CophosK+ provides reproducible predictions

To investigate whether the KSA predictions provided by CoPhosK^+^ are reproducible, we compared the prediction results from datasets I and II with the other two independent public MS-based phosphoproteomics datasets from human ovarian tumors and breast cancer xenograft tissue (IV and V). 543 phosphosites appear in both ovarian cancer datasets. 373 out of 543 phosphosites have a predicted kinase in KinomeXplorer and consequently in CophosK+. We have run CophosK+ on these new datasets and crosschecked the KSA predictions. Our results showed that the top predicted kinase of 155 phosphosites in this new ovarian cancer data is identical to the predicted kinase in the previous ovarian cancer data (i.e. 155/373 = 41% reproducibility rate for the top-ranked prediction). Moreover, the top-ranked kinase for 349 of the 373 phosphosites in the previous ovarian cancer dataset are ranked in the top 5 predicted kinases in this new dataset (i.e. 349/373 = 93% reproducibility rate for top-1 vs. top-5). There were 1899 common phosphosites between the two breast cancer datasets. CophosK+ has kinase predictions for 1079 out of 1899 phosphosites. Our result showed that, for these datasets, there are 22% and 40% reproducibility in top 1 and top 5 predicted KSAs, respectively.

### Enhancing KSA prediction by combining datasets

In Figure 7(a), we examine the overlap of CophosK^+^ based KSA predictions in the two biological contexts we consider (breast cancer and ovarian cancer). For this analysis, considering the top-ranked kinase as the prediction of CoPhosK^+^, we cluster the predictions by CophosK^+^ into three categories: 1. Predictions of CophosK^+^ that are consistent with those reported in PhosphoSitePlus. These include 402 predictions with 80 in common between breast and ovarian cancer datasets. 2. Prediction of CophosK^+^ is different from the kinase reported in PhosphoSitePlus. These include 393 predictions, with 29 in common. 3. No kinase annotation is available for that phosphosite in PhosphoSitePlus. These include 6425 phosphosites that are newly annotated with respect to a predicted KSA in this CophosK^+^ analysis, with 678 in common. Note that the predictions in the second category do not necessarily represent false positives, since a phosphosite can be targeted by multiple kinases and the annotations provided by PhosphoSitePlus are limited. The top-ranked kinase according to CophosK^+^ is identical for 109 phosphosites in categories 1 and 2. Among these, the kinase that is reported in PhosphoSitePlus is identical to that reported by CophosK^+^ for 80 phosphosites. If we define precision as the number of target kinases that are ranked first (category 1) divided by the total number phosphosites with a known target kinases in PhosphoSitePlus (union of category 1 and category 2), we can expect that (at least) 73% of top predictions which are identical between two different datasets will be correct. This improved accuracy also may be applicable in the context of expanded coverage for the 678 phosphosites that do not have any annotation in PhosphoSitePlus (category 3).

**Figure 7.**
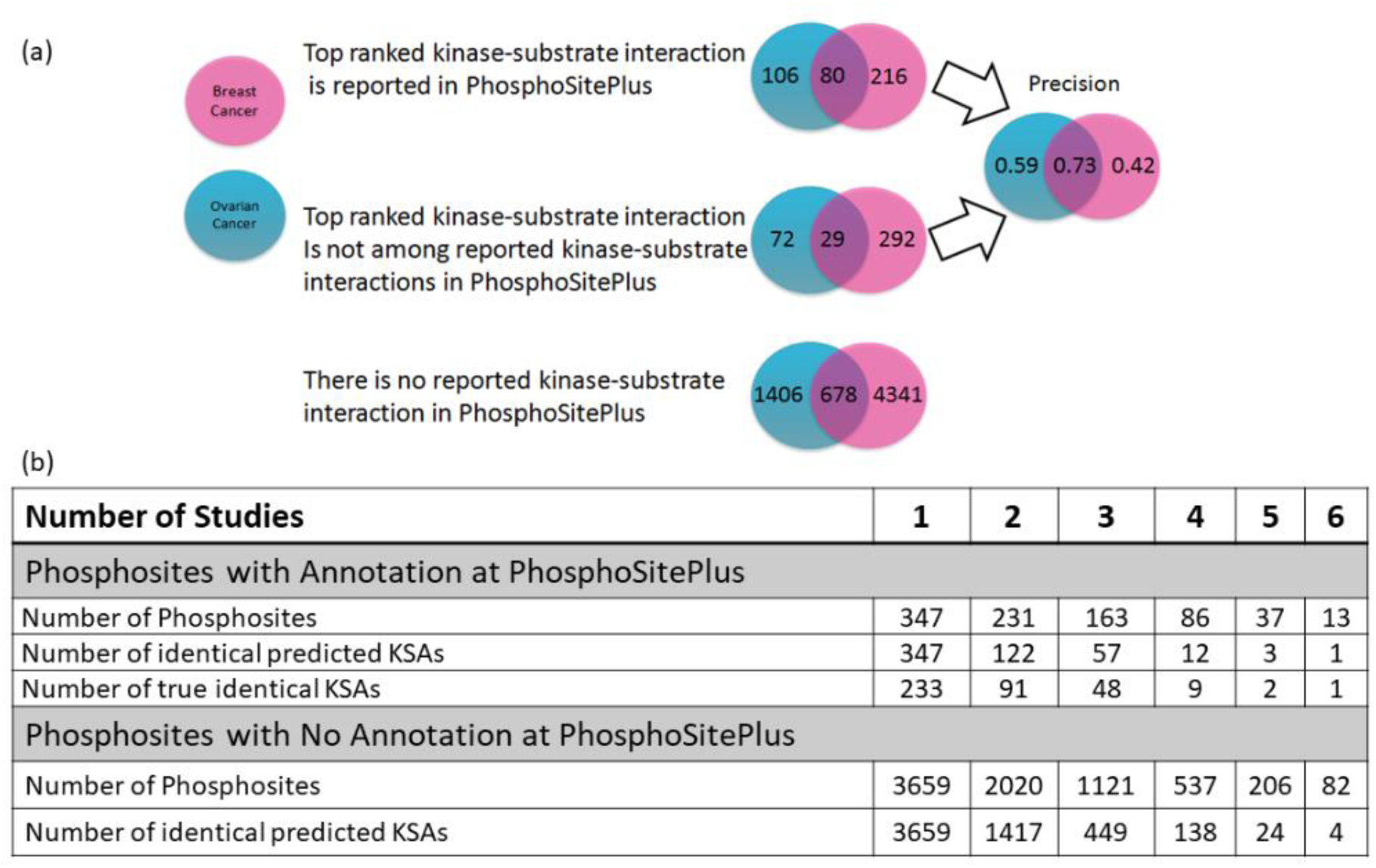
Consistency and reproducibility of kinase-substrate predictions made using different phosphorylation data sets. For each phosphosite in each data set, we rank the kinases using CophosK^+^, and then we identify the top ranked kinase based on two different datasets. **(a)** The Venn diagrams show (1) the number of phosphosites for which top-ranked kinase agrees with that reported in PhosphoSitePlus (True Positive), (2) the number of phosphosites that have kinases reported in PhosphoSitePlus but top-ranked kinase does not agree with that reported in PhosphoSitePlus (False Positive), and (3) the number of phosphosites with no kinase annotation reported in PhosphoSitePlus. The blue circles represent predictions based on ovarian cancer tumor cell lines, pink circles represents predictions based on breast cancer PDX. For the phosphosites for which at least one kinase is listed in PhosphoSitePlus, the right panel shows the “precision” (True Positive / (True Positive + False Positive)) of the top ranked kinase for each individual dataset and the intersection between the two datasets (i.e., the same kinase is ranked top in both datasets). **(b)** The number of phosphosites with at least one annotation in PhosphoSitePlus (upper panel) and no annotation in PhosphoSitePlus (lower panel) are shown as a function of the number of datasets that contain the phosphosite. Among these phosphosites, the number of phosphosites for which the top ranked kinase is identical across multiple datasets (identical predicted KSAs) is also shown as a function of the number of supporting datasets. For the phosphosites with annotations, the number of predictions that are consistent with PhosphositePlus annotations are also shown (true identical KSAs).

Motivated by the observation that predictions that are supported by two datasets have improved accuracy as compared to predictions that are supported by a single dataset, we also investigate the effect of the number of datasets supporting a prediction on the accuracy of that prediction. To characterize the trade-off between accuracy and coverage in integrating multiple datasets, we also assessed the number of predictions that are supported by multiple datasets as a function of the number of datasets. The results of this analysis are tabulated in Figure 7(b). As seen on the table, if the kinase that is ranked top by CophosK+ is identical across multiple datasets, it is more likely to be the kinase that is also reported in PSP, as compared to a candidate KSA that is predicted on only one dataset. Thus the accuracy is much improved when multiple phosphoproteomics datasets are available as compared to the accuracy of predictions provided by CoPhosK+ on a single dataset. These predictions are also drastically more accurate than the predictions of KinomeXplorer alone. While CoPhosK+ can provide predictions for thousands of sites using one or two datasets (e.g., the number of sites that are shared between two datasets is 2020 and the top-ranked kinase for 1417 of these 2020 sites are identical for the two datasets; only 122 of these 2020 have an annotation in PSP and the predicted kinase for 91 of these 122 sites is consistent with the PSP annotation.), there are only a few sites for which the predicted kinase is supported by at least five datasets. Nevertheless, as seen on the table, CoPhosK+ is able to provide predictions for more than 100 sites for which the predicted kinase is supported by four datasets (12 of which have annotations in PSP and 9 of these annotations are consistent with CoPhosK+’s predictions) and more than 400 sites for which the predicted kinase is supported by three datasets (57 of which have annotations in PSP and 48 of these annotations are consistent with CoPhosK+’s predictions).

### CophosK provides context-specific predictions of co-phosphorylation networks

As some phosphosites can be phosphorylated by multiple kinases, ascertaining which individual kinase is activated in different cell or tissues types is challenging to predict and yet crucial to the drug discovery process. Therefore, an important benefit of integrating MS-based phosphoproteomic data into KSA predictions is that these data can capture the context-specificity of these interactions. Clearly, no other method can consider a similar global biological context while predicting KSAs, since sequence motifs, structural information, and protein interactions considered by these methods do not represent a specific biological context, at least with the current state of cell based information. To investigate how CophosK^+^ captures context-specificity of KSAs, we identify common phosphosites between two datasets such that the kinase ranked as the top kinase by CophosK^+^ is different for different datasets, but all the kinase annotations are reported in PhosphoSitePlus (S1 Table). For example, NDRG1, the N-Myc downstream regulated 1, is known in PhosphoSitePlus to have multiple possible kinases, SGK1 and PRKACA, for which its phosphosite S330 represents a substrate target. However, a previous study shows that several Akt-inhibitor-sensitive breast cancer cells showed marked NDRG1 phosphorylation despite the low or undetectable level of SGK1 protein [30]. Co-phosphorylation analysis suggests strong correlation between the behaviour of SGK1 substrates and S330 in the ovarian cancer cell line while PRKACA annotated substrates track S330 more closely in the breast cancer models (S1 Table). NDRG1 is clearly annotated as having multiple and wide ranging roles in cellular stress response [31], proliferation and growth arrest [32], and tumor progression and metastasis [33] and a static prediction for its regulatory circuitry may not suffice for explaining its diverse roles. Our co-phosphorylation approach provides clear and testable predictions to uncovering these relationships in the relevant cellular or disease context. Nevertheless, they must be validated by knocking out the target kinase and assessing the effect on phosphorylation of specific sites of interest.

## Discussion and Future Work

In this paper, we present an integrative approach for KSA prediction using correlations among phosphosite intensities from phosphoproteomics data alone or coupled to sequence and protein interaction data to provide global and accurate predictions of KSAs. Although these advances are considerable, there is still much work to be done to better understand the power of co-phosphorylation analysis. For example, our method is limited by the coverage of phosphosites identified in LC/MS studies and prior knowledge of KSAs. The former limits the overlap between datasets making comparisons difficult while the latter limits the benchmarking of the approach. As our understanding of these phosphorylation overall improves, the tools will become more valuable over time. In particular, the functional meaning of negative correlations in the data needs further exploration.

Negative co-phosphorylation between a pair of phosphosites might occur for different reasons. One possibility involves sites on the same protein, where the phosphorylation of one site inhibits the phosphorylation of another site [28,34] thus exhibiting negative (and statistically significant) co-phosphorylation values. For negative correlations of sites on different proteins many explanations are possible, and should be considered. First, phosphorylation of a kinase may activate or inhibit the kinase; in the former case the kinases’ substrates will tend towards positive co-phosphorylation with that regulatory site on the kinase or negative co-phosphorylation when the effect of the phosphorylation is inhibitory [35]. Furthermore, it is expected that phosphatases regulated by phosphorylation may exhibit negative or positive co-phosphorylation with their substrates depending on whether the phosphorylation is activating or inhibitory towards the phosphatase. More subtle effects are also clearly possible, where a chain of kinases and phosphatases (essentially a pathway) may activate or deactivate a key regulatory node at the intersection of other regulatory circuits. Thus, negative or positive correlation “signals” can be generated across the cell resulting in the complex set of in interactions implied by this analysis.

In the context of gene co-expression analysis, partial correlation is often utilized to remove indirect effects of genes on each other, thereby revealing direct interactions [36]. The application of partial correlation in the construction of co-phosphorylation networks can also improve the accuracy of these networks, and thus can improve the accuracy of CoPhosK’s predictions. The application of partial correlation to the assessment of co-phosphorylation requires consideration of the relationships between the phosphorylation sites on the same protein, as well as the relationship between the expression of proteins and the phosphorylation levels of the sites on these proteins. The promising results presented in this paper, along with the availability of multi-omic data that includes measurements of protein expression and phosphorylation, pave the way for the application of such advanced statistical measures to co-phosphorylation analysis as well.

As functional signalling networks rely on many types of post-translational modifications (PTMs), an integrated correlation analysis framework for multiple PTMs must ultimately be developed to explain phenotype and may be effective in defining relationships between types of PTMs. Nevertheless, CophosK provides a strong data-driven approach for prediction of KSAs and drastically increases the coverage of the phosphosites for which kinase associations can be predicted. Thus, generation of more high-throughput MS-based phosphoproteomics data representing a variety of biological contexts can be used in conjunction with CophosK to better enable drug discovery and provide a deeper understanding of biological signalling.

## Methods

### Co-phosphorylation

Co-phosphorylation between phosphorylation sites is computed using Biweight midcorrelation. In contrast to other measures of correlation that use the mean to standardize observations, biweight midcorrelation uses the median. Biweight midcorrelation of two vectors *x* ∈ *R*^1×*m*^ and *y* ∈ *R*^1×*m*^ is computed as

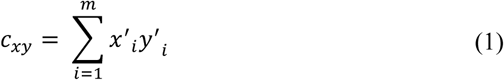

where,

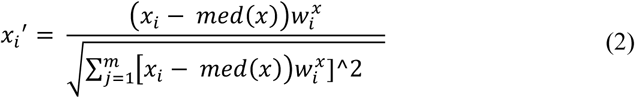

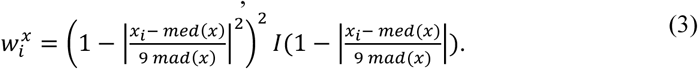

med(x) represents the median of vector x and mad(x) represents the median absolute deviation of vector x. I(u) is the indicator function which takes on value 1 if u > 0 and 0 otherwise. 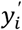 *and* 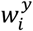 are also computed similarly for vector y.

### Model development for kinase prioritization

In order to prioritize the kinases for phosphorylation sites, we apply Bayes’ rule to derive the score for every pair of kinase-substrate.

#### Construction of co-phosphorylation network

Let *T* denote a set of all phosphorylation sites in PhosphoSitePlus and *P* denote the set of all phosphorylation sites that are present in the experimental data. We create a complete graph *G* in which the nodes represent the phosphosites in *P*. The weights of each edge are computed as the co-phosphorylation between the two corresponding phosphosites. Namely, for phosphosites *p, q* ∈ *P,* we denote the co-phosphorylation of *p* and *q* as *c*_*pq*_.

#### Scoring Schema (CophosK)

Note that it is possible to compute the co-phosphorylation of two phosphosites only if both phosphosites are present in the MS data (i.e, both sites are in *P*). Let *A* denote the distribution of co-phosphorylation among all pairs of phosphosites in *P* (i.e., *A* is the set of *c*_*pq*_ values across all (*p, q*) ∈ *P* × *P*). On the other hand, kinase information is available for the phosphosites that are in *T.* We call two phosphosites a *shared-kinase pair* if the two phosphosites are annotated as being regulated by the same kinase in PhosphoSitePlus. We denote the distribution of co-phosphorylation among shared-kinase pairs as *S* (i.e., *S* is the set of *c*_*pq*_ values across all (*p, q*) ∈ (*T* ∩ *P*) × (*T* ∩ *P*)).

For a kinase *k*, we define *T*_*k*_ ⊂ *T* as the set of phosphosites that are reported in PhosphoSitePlus as the substrates of kinase For a pair of phosphosites (*p, q*) ∈ *P*, Pr(*C*_*pq*_ > *c*_*pq*_|*S*) is the probability that the co-phosphorylation of *p* and *q* would be higher than *c*_*pq*_ given that *p* and *q* share a kinase. On the other hand, Pr(*C*_*pq*_ > *c*_*pq*_|*A*) represents the probability that the co-phosphorylation of *p* and *q* would be higher than *c*_*pq*_ for any pair of phosphosites. Using the Bayesian rule, we compute the log-likelihood of the association of phosphosite *p* with kinase *k* using the weights of edges between *p* and its neighbours that are in *Tk* :

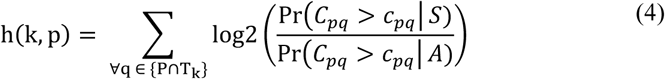

For a given phosphosite *p* ∈ *P,* CoPhosK computes all *h(k,p)* values (i.e. the log likelihood of association of kinase k and phosphosite p) for all kinases *k* and ranks the kinases in decreasing order of *h(k,p),* where a larger value of *h(k,p)* indicates that *k* is more likely to be a kinase that phosphorylates *p*.

#### Integrated Score (CophosK^+^)

To integrate the co-phosphorylation-based scores with static information, we have downloaded the pre-computed data for all available predictors on known phosphorylation sites from KinomeXporer-DB.version59. The scores reported by KinomeXplorer-DB represent the likelihood of association of kinases and substrates based on the integration of protein interaction based scoring and sequence-based scoring. Assume that the KinomeXplorer score for the interaction between kinase k to phosphosite p is *x(k,p)*. We compute CophosK^+^ score (i.e. *M(k,p))* for phosphosite p and kinase k by combining CophosK score and KinomeXplorer score as follows:

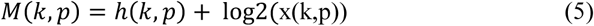

As in CoPhosK, CoPhosK+ also computes *M(k,p)* for all phosphosite-kinase pairs, and ranks kinases for each phosphosites such that a larger value of *M(k,p)* indicates that *k* is more likely to be a kinase that phosphorylates *p*.

#### Running PUEL

We downloaded the jar file provided byYang et al. and used the default parameters as initial parameters (Size of ensemble = 50, kernel type = radial). For each kinase, we run PUEL on our data using the known kinase-substrate interactions downloaded from PhosphoSitePlus as the training set. Then, for each phosphosite, we rank the kinases using the computed scores.

