## Supplementary Materials for "CoPhosK: A Method for Comprehensive Kinase Substrate Annotation Using Co-phosphorylation Analysis"

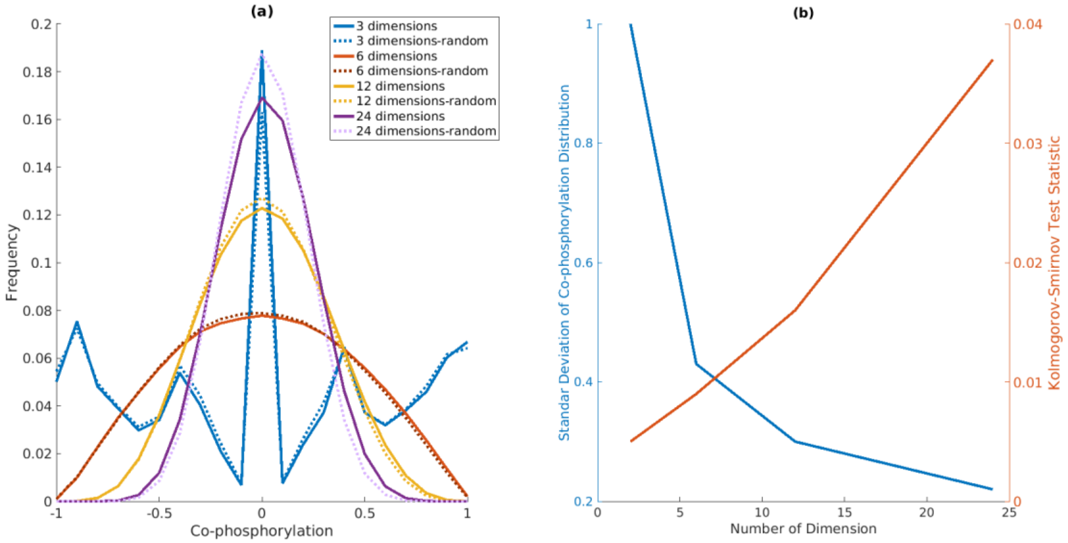


**S1 Fig. Effect of number of dimensions on co-phosphorylation distribution among phosphosite pairs in breast cancer PDX dataset.** Using the 24 breast cancer PDX samples, (a) we randomly selected subsets of samples and plot the co-phosphorylation distribution among phosphosites (solid line). We also randomize that data to compare against the co-phosphorylation distribution in original data (dashed line). (b) The standard deviation of the co-phosphorylation distribution is shown in the blue line and red line shows changes of Kolmogorov-Smirnov test statistic (the maximum absolute difference between cumulative distribution of co-phosphorylation in original and permuted data) in different number of dimensions.

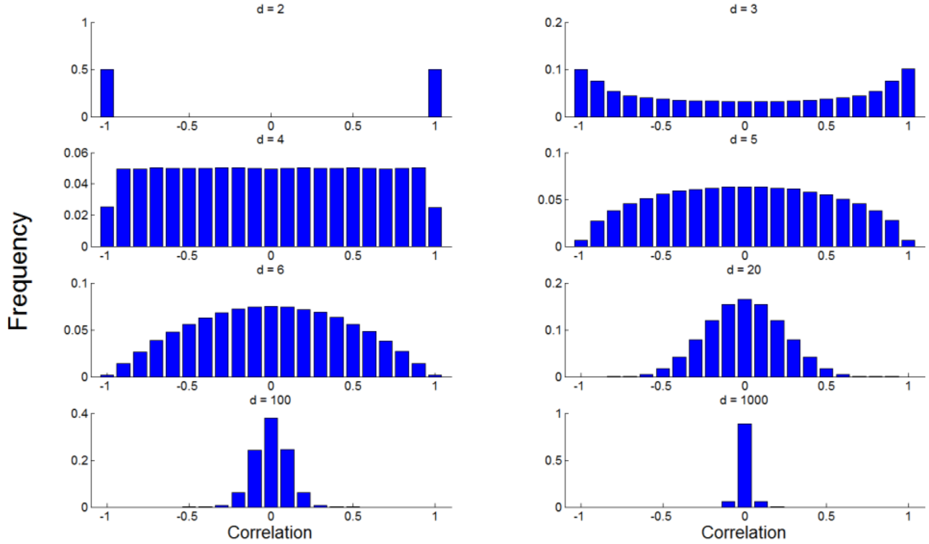


**S2 Fig. Correlation distribution among pairs of randomly generated vectors as a function of number of dimensions.** We generated 1000 random vectors from a normal distribution with zero mean and a standard deviation of one and plotted the distribution of correlation among all pairs of vectors. Each panel shows the histogram of correlations among pairs of vectors for a specific number of dimensions (denoted d).

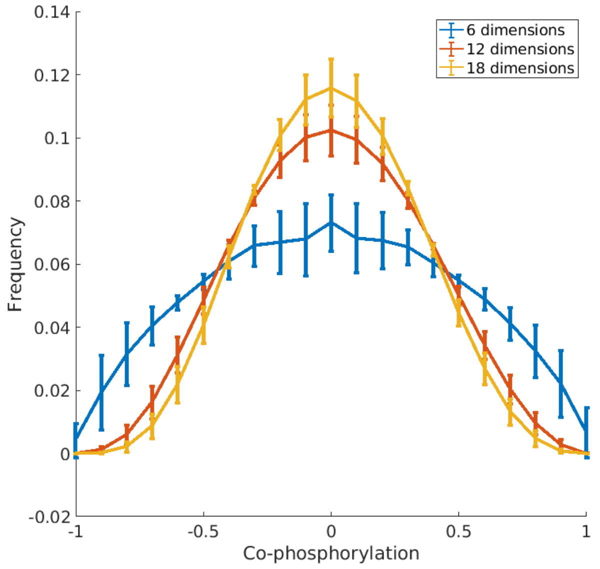


**S3 Fig. Effect of different sample selections on co-phosphorylation distribution among phosphosite pairs in breast cancer PDX dataset.** Using the 24 breast cancer PDX samples, we randomly select 6, 12 and 18 subsets of samples 100 times and plot the co-phosphorylation distribution among phosphosites.

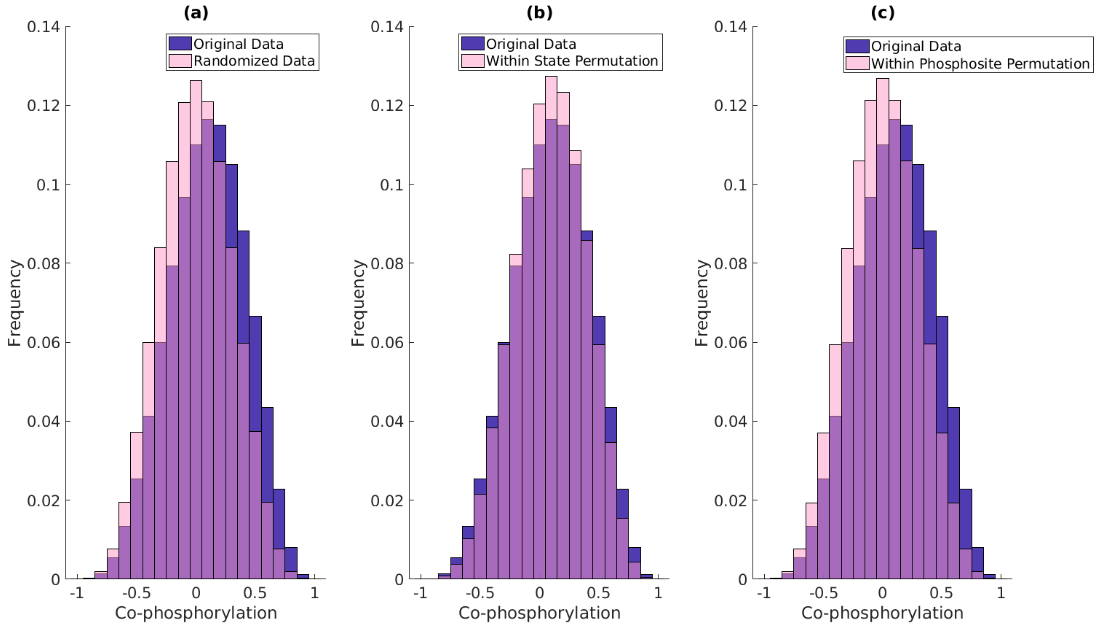


**S4 Fig. Co-phosphorylation distribution among phosphosite pairs in ovarian cancer tumors.** The blue histogram shows the distribution of co-phosphorylation (the correlation between the phosphorylation levels) of all pairs of phosphosites in ovarian cancer ($\mu$=0.09,$\sigma$=0.31). The pink histogram in each panel shows the distribution of co-phosphorylation of all pairs of phosphosites in 100 permutation tests representing (a) randomization of all entries in the phosphorylation matrix ($\mu$=1.6E-5,$\sigma$=0.29), (b) permutation of all entries across phosphosites for each state ($\mu$=0.08,$\sigma$=0.29), and (c) permutation of all entries across states within each phosphosite ($\mu$=1.3E-4,$\sigma$=0.29). The distribution of co-phosphorylation in the original dataset is significantly different as compared to the distribution of co-phosphorylation in all permutations (Kolmogorov-Smirnov (KS) test p-value << 1E-9).

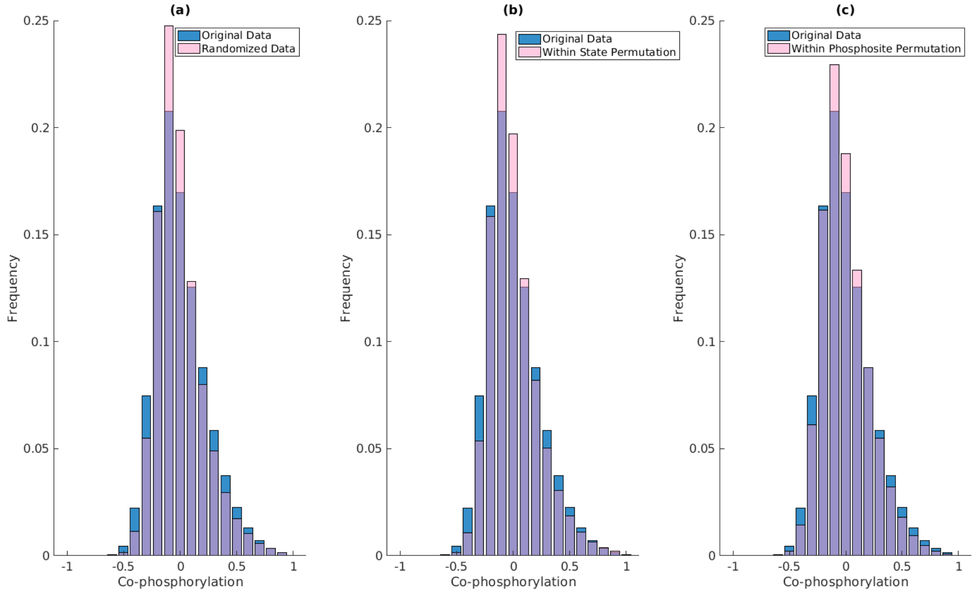


**S5 Fig. Co-phosphorylation distribution among phosphosite pairs in breast cancer PDX using Pearson correlation.** The blue histogram shows the distribution of co-phosphorylation (the correlation between the phosphorylation levels) of all pairs of phosphosites in breast cancer (μ=0.003,σ=0.23). The pink histogram in each panel shows the distribution of co-phosphorylation of all pairs of phosphosites in 100 permutation tests representing (a) randomization of all entries in the phosphorylation matrix (μ=-1.6E-5,σ=0.2), (b) permutation of all entries across phosphosites for each state (μ=0.005,σ=0.21), and (c) permutation of all entries across states within each phosphosite (μ=-1.9E-6,σ=0.2). The distribution of co-phosphorylation in the original dataset is significantly different as compared to the distribution of co-phosphorylation in all permutations (Kolmogorov-Smirnov (KS) test p-value << 1E-9).

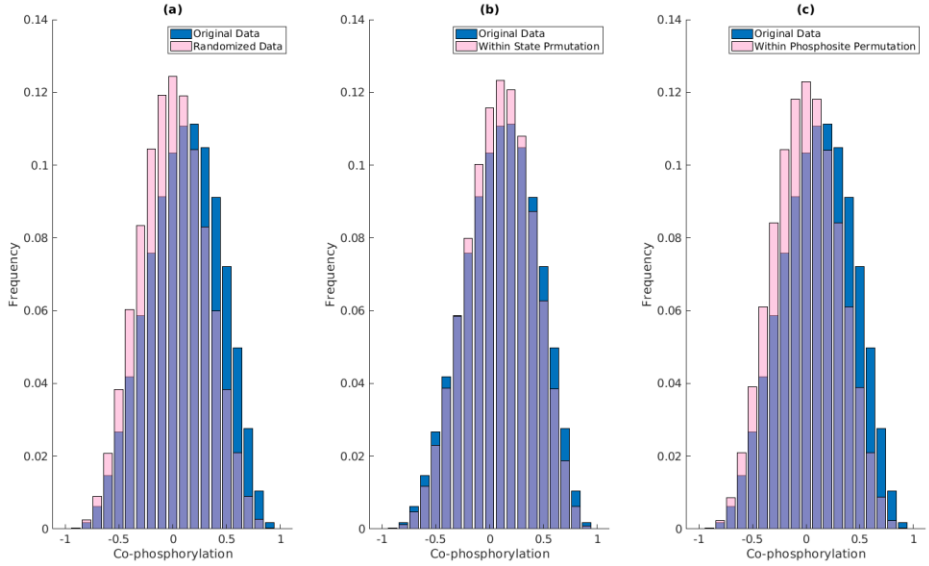


**S6 Fig. Co-phosphorylation distribution among phosphosite pairs in ovarian cancer tumors using Pearson correlation.** The blue histogram shows the distribution of co-phosphorylation (the correlation between the phosphorylation levels) of all pairs of phosphosites in breast cancer (μ=0.1,σ=0.32). The pink histogram in each panel shows the distribution of co-phosphorylation of all pairs of phosphosites in 100 permutation tests representing (a) randomization of all entries in the phosphorylation matrix (μ=-2.6E-5,σ=0.3), (b) permutation of all entries across phosphosites for each state (μ=0.09,σ=0.3), and (c) permutation of all entries across states within each phosphosite (μ=-3.4E-5,σ=0.3). The distribution of co-phosphorylation in the original dataset is significantly different as compared to the distribution of co-phosphorylation in all permutations (Kolmogorov-Smirnov (KS) test p-value << 1E-9).

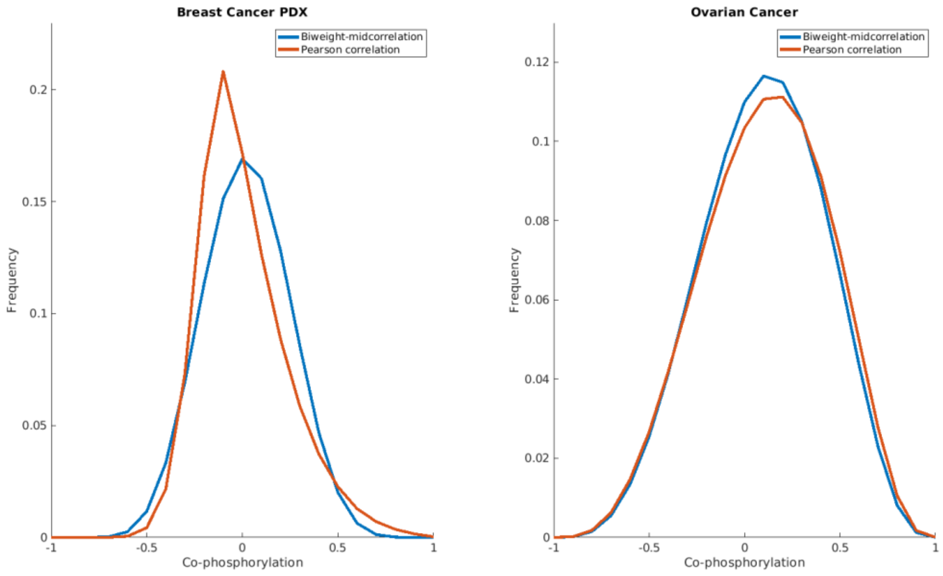


**S7 Fig. Comparison of co-phosphorylation distribution among phosphosite pairs in (a) breast cancer PDX and (b) ovarian cancer tumors using biweight-midcorrelation (blue curve) and Pearson correlation (red curve).**

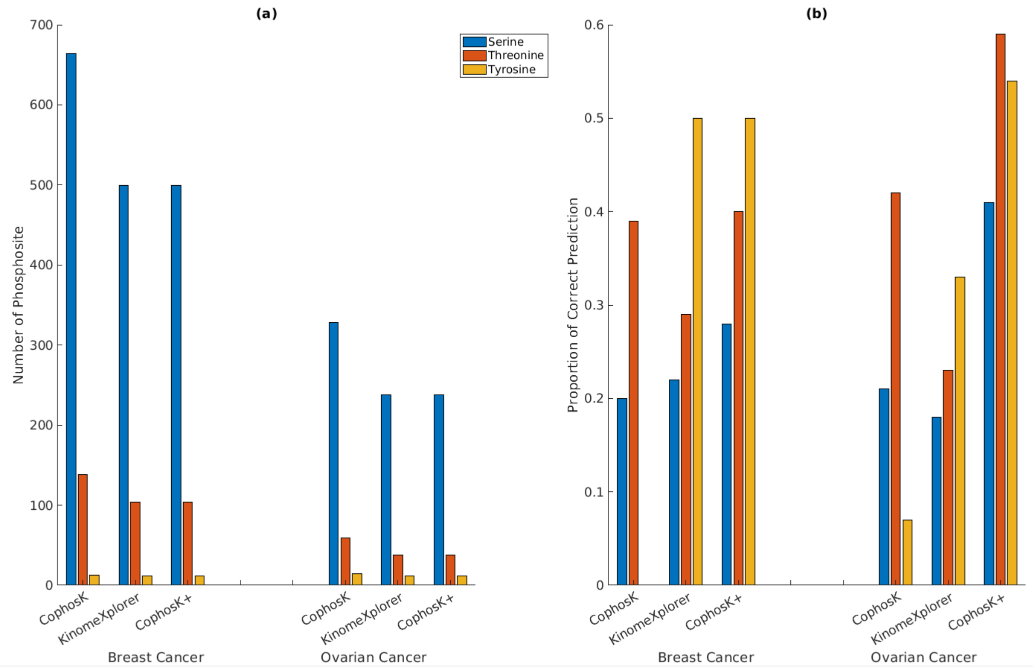


**S8 Fig. Number of phosphosite and site-specific prediction performance.** Number of annotated phosphosites and the methods' performance separated on specific residue is reported in (a) and (b), respectively.

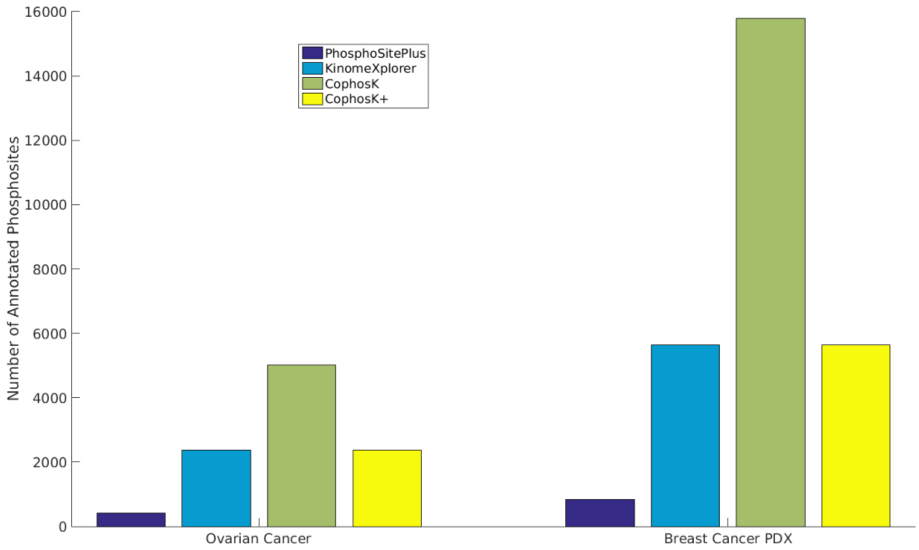


**S9 Fig. Coverage of kinase-substrate interaction predictions.** Number of phosphosites in each dataset that are annotated by each method is shown.

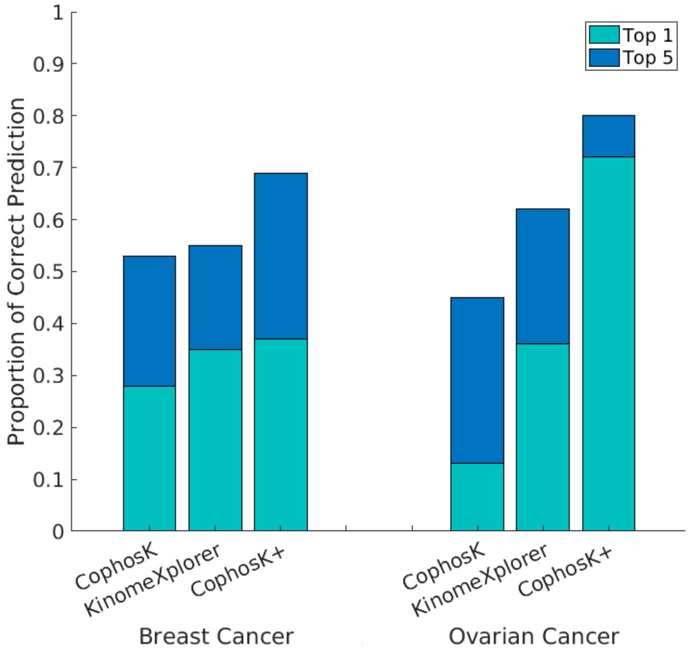


**S10 Fig. Performance of CophosK, KinomeXplorer and CophosK+ in predicting kinases for phosphosites in breast cancer and ovarian cancer data using phosphor.ELM as predefined KSAs.** There is 2427 KSAs reported in the Phospho.ELM dataset and 1350 KSAs are common between PhosphoSitePlus and Phospho.ELM

**S1 Table. Data-specific kinase prediction.** The phosphosites listed in this table are reported to have more than one kinase in PhosphoSitePlus. CophosK$^{+}$ identifies previously reported, but different kinases as the top-ranked candidate based on each dataset.

| Protein | Site | Reported Kinase in PhosphoSitePLUS | KinomeXplorer prediction | CophosK+ prediction on ovarian cancer | CophosK+ prediction on breast cancer |
|---|---|---|---|---|---|
| BAD | S99 | AKT1,BRAF,PAK1,PRKACA | AKT1 | AKT1 | PRKACA |
| FLNA | S2152 | PAK1,PRKACA,AKT1,RPSGKA | AKT1 | PAK1 | PRKACA |
| HIST1H1E | T18 | CDK1,CDK2 | DAPK3 | CDK1 | CDK2 |
| NDRG1 | S330 | SGK1,PRKACA | SGK1 | SGK1 | PRKACA |
| PFKFB2 | S466 | AKT1,PRKACA,AMPKA1 | AKT1 | PRKACA | AKT1 |
